# Complete fold annotation of the human proteome using a novel structural feature space

**DOI:** 10.1101/092379

**Authors:** Sarah A. Middleton, Joseph Illuminati, Junhyong Kim

**Affiliations:** Genomics and Computational Biology Graduate Program, University of Pennsylvania, Philadelphia, PA 19104, USA; Department of Computer Science, University of Pennsylvania, Philadelphia, PA 19104, USA; Department of Biology, University of Pennsylvania, Philadelphia, PA 19104, USA

## Abstract

Recognition of protein structural fold is the starting point for many structure prediction tools and protein function inference. Fold prediction is computationally demanding and recognizing novel folds is difficult such that the majority of proteins have not been annotated for fold classification. Here we describe a new machine learning approach using a novel feature space that can be used for accurate recognition of all 1,221 currently known folds and inference of unknown novel folds. We show that our method achieves better than 96% accuracy even when many folds have only one training example. We demonstrate the utility of this method by predicting the folds of 34,330 human protein domains and showing that these predictions can yield useful insights into biological function, including the prediction of functional motifs relevant to human diseases. Our method can be applied to *de novo* fold prediction of entire proteomes and identify candidate novel fold families.

## Introduction

Although protein sequences can theoretically form a vast range of structures, the number of distinct three-dimensional topologies (“folds”) actually observed in nature appears to be both finite and relatively small^1^: 1,221 folds are currently recognized in the SCOPe (Structural Classification of Proteins—extended) database^2^, and the rate of new fold discoveries has diminished greatly over the past two decades. Nevertheless, extending the catalog of protein fold diversity is still an important problem and fold classifying the entire proteome of an organism can lead to important insights about protein function^3–5^. Large-scale fold prediction typically involves computational methods, and the computational difficulty of *ab initio* structure prediction has led to template matching (e.g., using methods such as HHPred^6^) as the most common method for predicting the structure. When sequence-based matching is difficult, other fold recognition approaches must be employed, such as protein threading. Threading-based methods, especially those that combine information from multiple templates, have been among the most successful algorithms in recent competitions for fold prediction^7,8^, but are bottlenecked by long run times. Machine learning-based methods have also been used, which can be designed either to recognize pairs of proteins with the same fold^9,10^ or classify a protein into a fold^11,12^. Although these methods have shown promising results for a subset of folds, they have so far not been able to generalize to the full-scale fold recognition problem. This failure can mainly be attributed to the severe lack of training data available for most SCOPe folds, as well as the highly multi-class nature of the full problem, which requires distinguishing between over 1,000 different folds^12^.

Here we introduce a method for full-scale fold recognition that integrates aspects of both threading and machine learning. At the core of our method is a novel feature space constructed by threading protein sequences against a relatively small set of structure templates. These templates act as “landmarks” against which other protein sequences can be compared to infer their location within structure space. We show the utility of this feature space in conjunction with both support vector machine (SVM) and first-nearest neighbor (1NN) classifiers, and further develop our 1NN classifier into a full-scale fold recognition pipeline that can predict all currently known folds. Applied to the entire human proteome, our method achieves 95.6% accuracy on domains with a known fold and makes thousands of additional high-confidence fold predictions for domains of unknown fold. We demonstrate utility by inferring new functional information, focusing on RNA-binding ability. The structure and function annotations of the entire human proteome are provided as a resource for the community.

## Results

### The protein empirical structure space (PESS)

Our approach is based on the idea of an empirical kernel^13^, where the distance between two objects is computed by comparing each object to a set of empirical examples or models. We have previously applied this idea to RNA secondary structure analysis^14^, and we show here that it can be adapted to proteins. The objects being compared are amino-acid sequences and the distance we would like to compute is similarity of tertiary structure. We selected a set of 1,814 empirical threading templates that describe the three-dimensional coordinates of atoms of proteins of known structures. We use only a small subset of known structures for our template library which we find sufficient to construct an informative structural distance function. Using the threading templates we mapped amino-acid sequences to a structural feature space, where the coordinates of each sequence reflect its threading scores against the templates. We refer to this as the protein empirical structure space (PESS). Using the PESS, we trained a classifier to recognize every fold (Fig. 1). Since protein domains are the unit of classification in SCOPe, we applied this approach to protein domains as units rather than full proteins.

**Figure 1.**
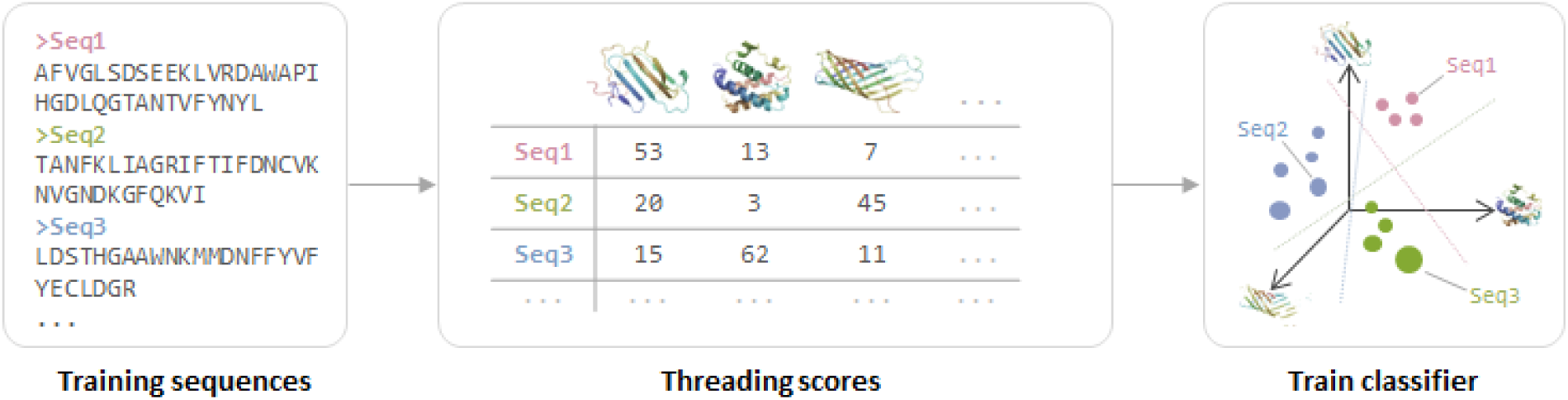
Overview of PESS construction. Training sequences of known fold are threaded against a set of structure templates, and the resulting threading scores act as coordinates within a structural feature space (the PESS). A classifier can then be trained to recognize the subspace occupied by each fold in the PESS. Different colors indicate the fold of each sequence and are shown here only for visualization.

### Fold recognition performance

We tested the PESS in combination with 1NN or SVM classifiers (Fig. 2a & b) using three popular benchmarks from the TAXFOLD paper^12^. These benchmarks are designed to test the ability of a method to distinguish between increasing numbers of folds: 27 folds in EDD, 95 in F95, and 194 in F194. Each fold has at least 11 training examples. The accuracy of our classifiers are shown in Table 1 along with several other published methods^12,15–20^. Our SVM classifier performed the best on all three benchmarks, with the exception of the EDD dataset, where the best performance was from the method of Zakeri et al. when it was used in combination with known Interpro functional annotations. Our 1NN classifier also performed very well on all three benchmarks, outperforming all but our SVM on F95 and F194. We next asked whether our method actually performed better than simply using the top-scoring template from our feature space. We found that directly using the fold of the top template resulted in 52.1, 56.4, and 57.4% accuracy on EDD, F95, and F194 respectively. Therefore, using the threading scores as a feature space rather than for direct classification improved performance considerably.

**Figure 2.**
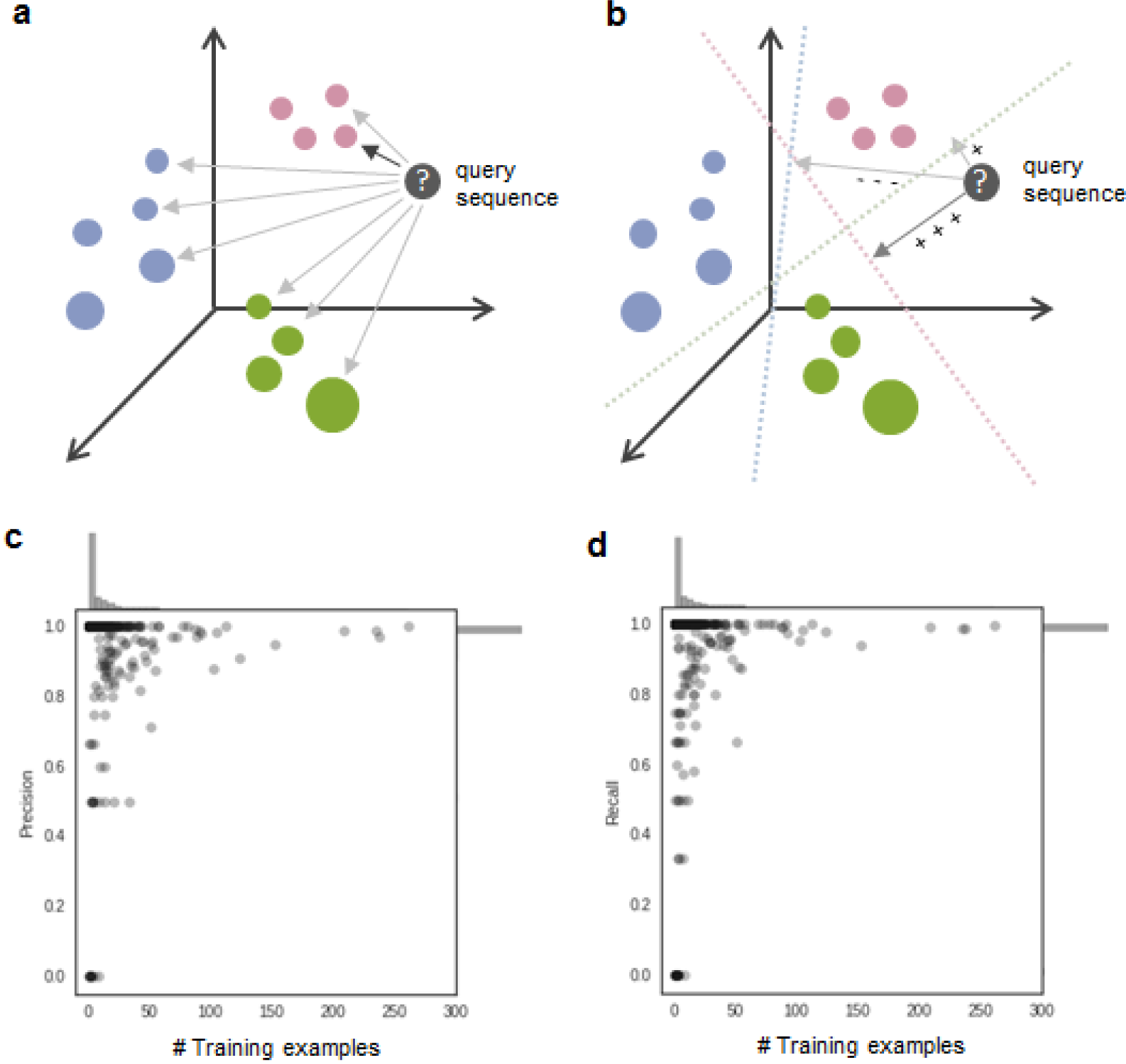
Classification and performance using the PESS. (a & b) Two different methods of classification using the PESS. Colored circles represent training examples within the PESS and are colored by fold. (a) In 1NN classification, the PESS distance between the query (gray circle) and all training examples is computed and the query is assigned to the fold of the nearest training example (dark gray arrow). (b) In 1-vs-all SVM classification, the PESS distance between the query and each of the fold-level hyperplanes (dotted lines) is computed, and the query is assigned to the fold that gives the best score (dark gray arrow), based on signed distance from the fold’s hyperplane. (c) Precision and (d) recall measures were computed for each fold separately after 1NN classification using the PESS and plotted against the number of training examples for each fold. Marginal histograms show the distribution of folds along each axis.

**Table 1.** Overall% accuracy on three benchmarks using 10-fold cross validation

| **Method** | **EDD** | **F95** | **F194** |
| --- | --- | --- | --- |
| Dehzangi et al.* | 88.2 | - | - |
| Saini et al.* | 86.6 | - | - |
| Zakeri et al. | 88.8 / 96.9** | - | - |
| HMMFold * | 93.8 | - | - |
| TAXFOLD | 90.0 | 82.4 | 79.6 |
| PFPA* | 92.6 | 83.6 | 78.2 |
| This method – 1NN | 90.6 | 84.6 | 82.5 |
| This method - SVM | 95.7 | 91.9 | 90.5 |

\* These papers used slightly modified versions of the datasets
\*\* With Interpro functional annotations

The benchmarks described above included only a subset of the 1,221 folds in SCOPe v.2.06. Recognizing all folds simultaneously is challenging; not only is it a highly multiclass problem, but it also suffers from a lack of training examples for a large fraction of the folds. We focused on our 1NN classifier, which requires only a single training example per fold, to scale to the full fold recognition task. To train the classifier to recognize all folds, we downloaded domain sequences from the Astral database^2^ corresponding to SCOPe (v.2.06) filtered to less than 20% pairwise identity, which we call SCOP-20. This dataset contains 7,659 sequences covering all 1,221 folds in classes “a” through “g”. To create a separate test set, we also downloaded the SCOPe sequences filtered to 40% identity and then removed any overlap between this set and the SCOP-20 set. This resulted in 6,322 sequences in 609 folds, which we call the SCOP-40 dataset. Using 1NN classification, 97.6% of SCOP-40 domains were classified into the correct fold (precision=0.964, recall=0.95). Using a combined SVM+1NN classifier (see Methods) did not improve performance (acc=96.9%, precision=0.917, recall=0.938), indicating that the 1NN classifier alone is sufficient for good classification on this dataset. Of the folds represented in the SCOP-20 training set, 86.5% (1,055) have fewer than 10 training examples, and almost half (605) are “orphan” folds with only one training example. Accurate classification into these folds is expected to be particularly difficult due to the small amount of training data. To determine how well our method performs relative to the number of training examples, we calculated precision and recall separately for each fold based on the SCOP-40 classification results. Although performance on folds with fewer training examples was slightly worse overall, the vast majority of folds had perfect precision and recall, regardless of training size (Fig. 2b & c). Focusing specifically on orphan folds, for which classification should be most difficult, we found that 96.4% of the 275 training examples belonging to these folds were correctly classified, which was only slightly lower than the overall SCOP-40 accuracy. Thus, our method can accurately recognize folds even when there is a single training example.

### Proteome-scale fold prediction of human proteins

The ability of the PESS to accurate recognize all folds with relatively little threading makes it well suited for classifying large, proteome-scale datasets. Here we applied our new method to predicting the fold of protein domains curated from the entire human proteome. Since the 1NN-only classifier performed better than the SVM+1NN combined classifier on the full-scale fold recognition test, we used the 1NN-only classifier to predict the folds of all human protein domains.

An overview of our whole proteome fold classification pipeline is shown in Figure 3a. In contrast to SCOP-derived benchmarks, whole proteomes present several additional challenges for fold recognition. One of the major bottlenecks is the process of segmenting whole proteins into domains, which is often slow and error-prone. We did not attempt to address this issue here, but instead make use of the existing domain segmentation of the human proteome performed by the Proteome Folding Project^5^. Another challenge is recognizing domains that do not belong in any of the known fold categories, e.g. due to segmentation errors, being disordered, or belonging to a previously undiscovered fold. To address this problem, we defined a distance threshold for classification based on the typical distance between a domain and its nearest neighbor when the true fold of the domain is not represented in the feature space (see Methods). When a query domain’s nearest neighbor is farther than this threshold distance, the domain is assigned to a “no classification” category (Fig. 3a).

**Figure 3.**
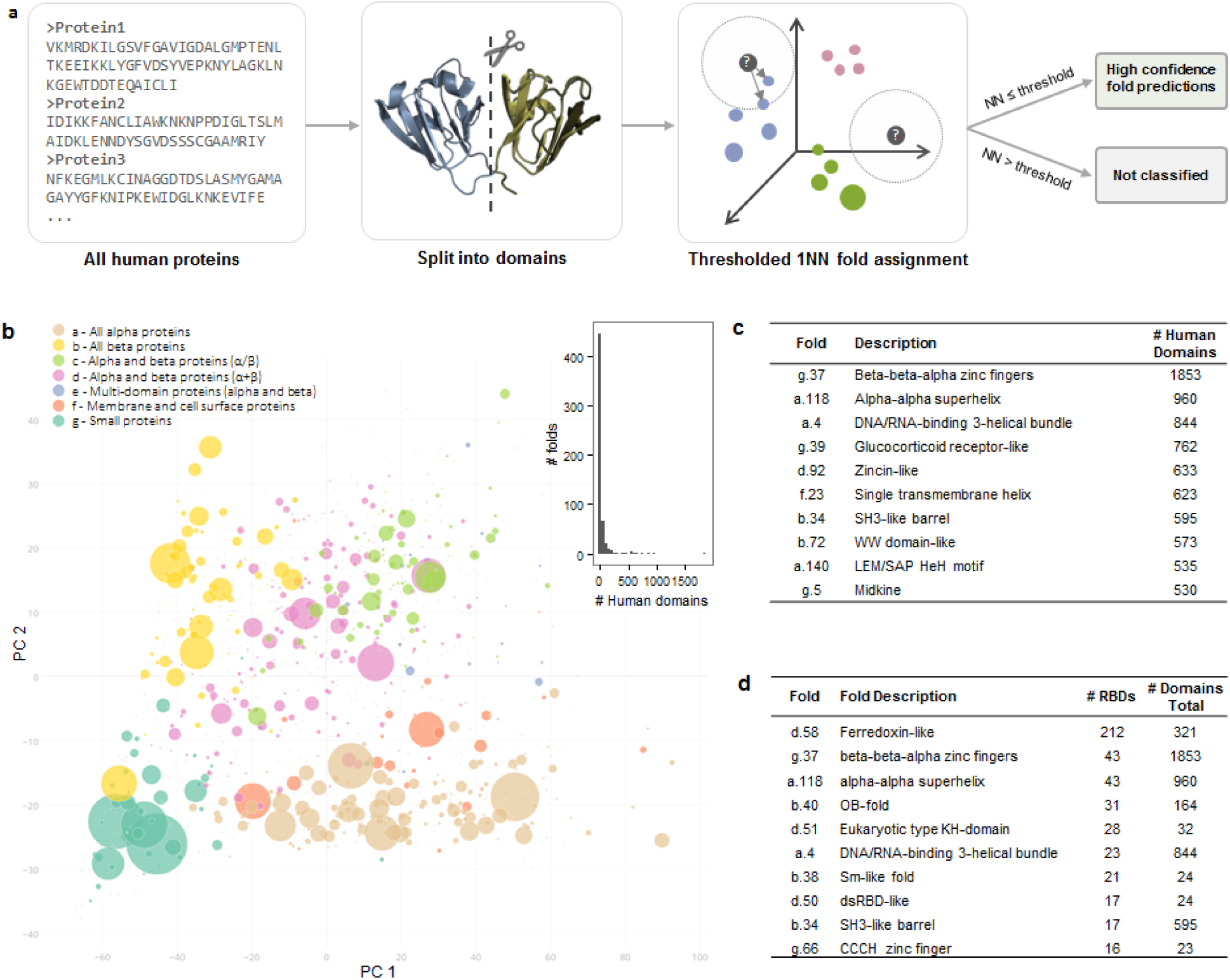
Fold classification of the human proteome. (a) Overview of classification process. Full length human protein sequences were split at predicted domain boundaries to create one or more separate domain sequences per protein (Drew et al. 2011). Domain sequences were mapped to the PESS and classified by 1NN classification. A threshold was applied to the nearest neighbor distance (dotted circle), whereby only domains with a nearest neighbor closer than the threshold distance were classified. (b) PCA projection of fold centroids within the PESS, scaled by number of human domains predicted to belong to that fold. Centroids were calculated based on the location of each fold’s training examples within the PESS and are colored by SCOP class. Inset: distribution of the number of human domains predicted per fold. (c) Top ten folds by number of human domain predictions. (d) Top ten likely RNA-binding folds, ranked by number of confirmed RNA-binding domains (RBDs). Confirmed RBDs were determined based on matches to a curated list of RNA-binding related Pfam families.

There were a total of 34,330 human domains with length greater than 30 residues in the Proteome Folding Project dataset, corresponding to 15,619 proteins. Of these, 20,340 domains (59%) had a nearest neighbor within the distance threshold and were classified into an existing fold by our method. Only 128 of these domains were previously placed into a fold with high confidence by the Proteome Folding Project^5^. To test how well our predictions match with what is currently known about human protein structures, we used a blastp search against PDB to identify 2,211 human domain sequences with a “known” fold; that is, an identical or highly similar PDB entry with a SCOPe fold classification. Our classifier made a fold prediction for 1,873 (84.7%) of these domains, and 95.6% of these predictions exactly matched the known SCOPe fold.

Overall, 757 of the 1,221 SCOPe folds had at least one human domain predicted by our method. The distribution of domains across folds was highly skewed, with the majority of folds having only a few predicted domains and a small number of folds having many (Fig. 3b). This agrees with previous observations that domains are not evenly distributed in protein structure space^1,21^. The top 10 folds accounted for 38.9% (7,908) of the classified domains, and the most common fold (Beta-beta-alpha zinc fingers) alone encompassed 9.1% (1,853) of the fold predictions (Fig. 3c). A full list of fold predictions is provided in Supplementary Table 1.

#### Human RNA-binding proteins

RNA-binding proteins (RBPs) are an important class of proteins that function in almost all aspects of RNA biology, including splicing, translation, localization, and degradation. It would be valuable to fully define which folds have potential RNA binding function and use this information to improve our annotations of RBPs. We obtained a list of 1,541 currently known RBPs in humans from a recent RBP census^22^ and extracted the corresponding domains from our dataset. There were 1,816 domains with fold predictions, matching 243 different folds.

Since not every domain in an RBP is expected to actually bind RNA, we first sorted these folds into “likely RNA-binding domain (likely RBD)” and “likely auxiliary” groups. The RBPs in the census were primarily identified based on hits to a list of Pfam families with RNA-binding function, so we defined the likely RBD folds as those with at least two RBP domains with a hit (E < 0.01) to this RNA-binding Pfam list. There were 720 such domains which encompassed 78 different folds. The most common folds included several with well characterized RNA-binding function, such as Ferredoxin-like, which includes the RNA recognition motif (RRM); Eukaryotic type KH-domain (KH-domain type I); and dsRBD-like (Fig. 3d). Next, we defined the auxiliary folds as those with at least one RBP domain but fewer than two hits to the RNA-binding Pfam list. By this criteria, we identified 165 folds, the most common being the Cytochrome C fold (14 domains) and RING/U-box E3 ligase fold (12 domains). These folds are likely to represent other functions performed by the RBPs; however, we note that the lack of a Pfam match does not preclude RNA-binding function, so some of these auxiliary folds may in fact be RNA-binding.

The RBP census contained 21 cases where a protein was known to bind RNA but the type of RBD was not yet identified. Using our method, we matched three of these RBPs to one or more of the likely-RBD folds established above. One of these RBPs was Fam120a (also called C9orf10), which was previously found to have RNA-binding activity at its C-terminal end, but the type of RNA binding domain was not determined^23^. Our method predicted a DNA/RNA-binding 3-helical bundle fold within the RNA-binding region of this protein. Loosening the classification threshold slightly (NN distance ≤ 20) allowed us to identify potential RBDs for three more of the RBPs, including a partial Ferredoxin-like fold at the N-terminal of Int8 and a PABP domain-like fold in Int10.

We next looked to see if there were any additional proteins represented in the likely-RBD folds that were not already annotated as being RBPs by the census. We found 6,249 such proteins, which is over four times the number of RBPs currently known. For many of these proteins, we cannot confidently predict their RBP status based on fold alone because some of the likely-RBD folds have other functions besides RNA-binding (e.g. some superfamilies of the Ferredoxin-like fold can be protein binding instead of RNA binding). Nonetheless, several of the likely-RBD folds appear to be highly enriched in known RNA-binding domains, suggesting that functional annotation transfer is possible for these folds. For example, of the 32 domains predicted by our method to have the KH-domain fold, only four did not have a hit to the RNA-binding Pfam list, and of these, three were already known to be KH-domain RBPs based on the RBP census. The one domain that was not in the census was part of the Blom7 protein (also called KIAA0907), which has an experimentally determined structure (PDB: 2YQR) that confirms structural similarity to the KH-domain, despite the lack of a Pfam match. A full list of our new RBP predictions and likely-RBD folds is in Supplementary Table 2.

#### Novel folds in the human proteome

Each year at least a few new folds are added to SCOPe (e.g. 13 new folds were added in the latest release). As noted above, there were ~14,000 human protein domains, or ~40% of domains, that were not assigned to known folds. While some of these might be due to problems of segmentation, we hypothesize many of them represent uncharacterized folds. As a preliminary analysis of potential novel folds in the human proteome, we extracted a set of human domains that were not close to any of our training examples (NN distance ≥ 30) and clustered them (Methods). This resulted in 36 clusters (Fig. 4a and Supplementary Table 3), which we examined for evidence of novel folds.

**Figure 4.**
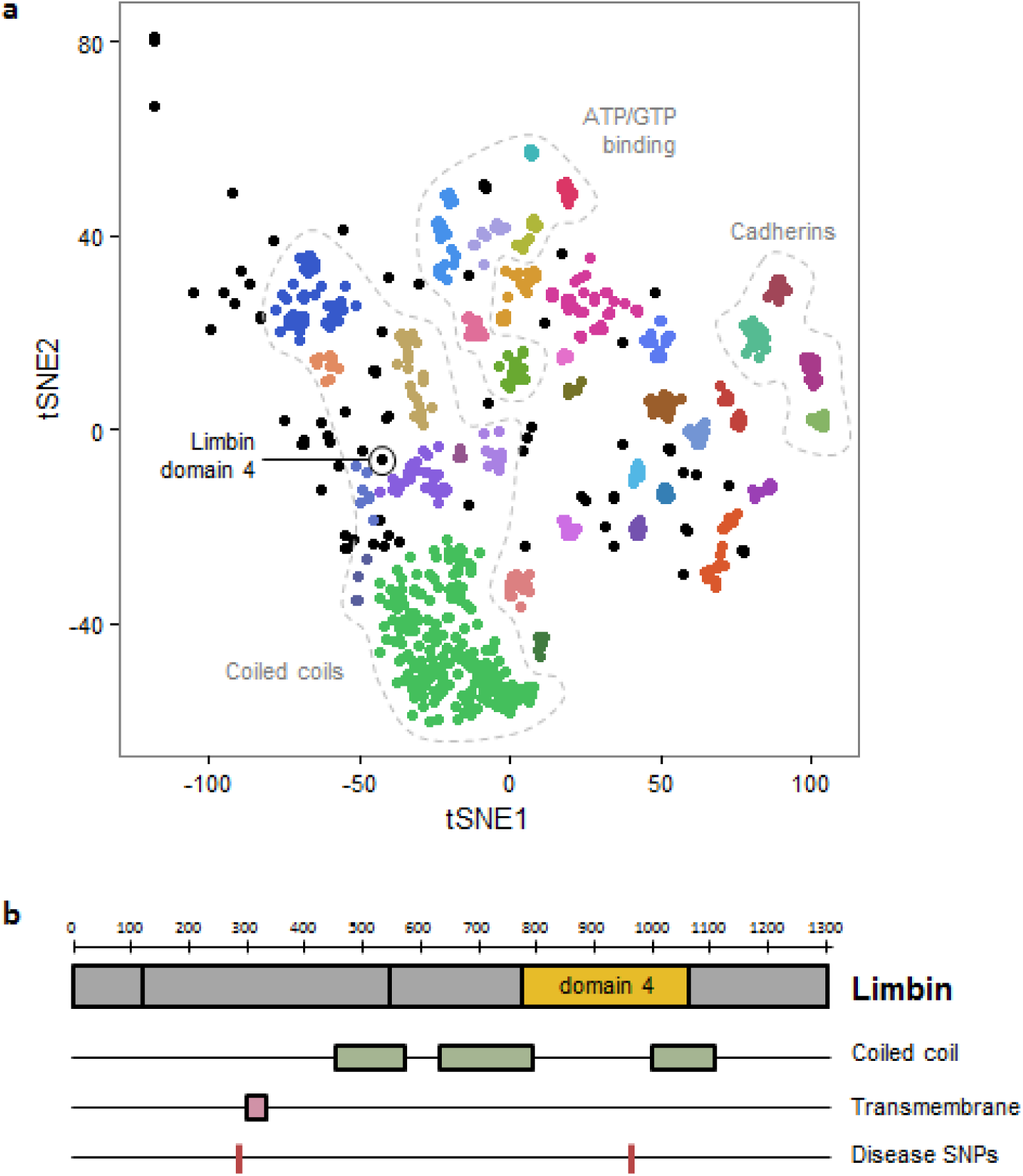
Analysis of unclassified human domains. (a) t-SNE projection of human domains with nearest-neighbor distance ≥ 30. Colors indicate cluster assignment by DBSCAN; unclustered domains are shown in black. Dotted lines show related groups of domains. (b) Overview of the *EVC2* protein product, Limbin, and its known structure elements. The location of the domain with a putative novel fold is shown in yellow.

We first looked for incorrect domain boundary prediction or errors of our prediction method. Many of the domains were unusually long (>500 residues) compared to the average domain in the training set (195 residues), suggesting that they may in fact be multiple domains. For example, there were four neighboring clusters that contained almost exclusively domains from the Cadherin family of proteins. Most of these domains were longer than 500 residues and overlapped multiple repeats of the Cadherin motif based on Pfam annotations. The Cadherin fold is modeled as a single repeat in SCOPe, so this is likely a case where fold classification failed due to improper domain definition. A similar problem was observed for six clusters containing domains from several different classes of ATP/GTP binding proteins, where each domain spanned multiple distinct Pfam annotations that are likely to represent separate folds. Overall, we found that 26 of the clusters were potentially the result of such segmentation errors.

The largest cluster contained 208 domains, most of which were of a reasonable length (289 residues on average). On closer examination, we found that a large fraction of these domains were predicted to have a coiled coil structure. The SCOPe hierarchy places most coiled coil domains in a separate class (class H) that was not included in the training data. Therefore, this cluster can possibly be explained by the absence of the correct fold within our training data, although it is not truly novel. Eight other neighboring clusters were also found to have predominantly coiled coil structure, indicating that these structures can potentially explain a substantial fraction of our unclassified domains.

We also examined the un-clustered domains, which might be isolated examples of novel folds. One domain, the fourth predicted domain of the protein Limbin (residues 775–1067), was found not to overlap any known Pfam, SCOP, or other structural annotation. Although this domain was located in the feature space in proximity to the coiled coil clusters (Fig. 4a), it is predicted to be only partially coiled coil (Fig. 4b). We performed a more thorough template search for this domain using HHPred^24^, RaptorX^25^, and SPARKS-X^26^ webservers, but did not identify a significant template match. Limbin is the protein product of the gene *EVC2*, which is involved in the hedgehog signaling pathway and is frequently mutated in Ellis-van Creveld syndrome^27,28^. Interestingly, one of the mutations linked to this disease is found within our domain of interest (Arg870Trp; rs137852928)^27^, suggesting that this region is functionally important. Whether this region represents a truly new fold will require additional analysis, but overall these results support the idea that the PESS can be used to identify novel structure groups.

## Discussion

Here we have demonstrated the utility of an empirically derived structural feature space composed of threading scores (the PESS) for addressing the problem of fold recognition. The most important characteristics of such a multi-dimensional feature space are the ability to combine characteristics of multiple fold templates for fold recognition and the ability to potentially identify entirely novel folds through interpolation of the feature space. Many types of classifiers can be used in conjunction with this feature space; we showed here that linear SVM achieved good performance on benchmarks where at least 10 training examples were available per fold, and 1NN worked well in the more general case to recognize all known folds. We applied our method to the human proteome, predicted high confidence fold classifications for 20,340 domains, and showed that these predictions can be used to make functional inferences as illustrated by the class of RNA-binding proteins. A distinct advantage of the PESS is that it only requires a single training example per fold when used in conjunction with a 1NN classifier, allowing us to make predictions for all currently known folds in SCOPe. This is critical, since almost half of all SCOPe folds have only one training example in SCOP-20. Another advantage of the 1NN classifier is that adding new training data does not require re-training the whole classifier, making it simple to update the model as new data become available.

One of the limitations of methods that rely on threading is the large amount of time the threading process takes. Threading against all PDB templates can take hours or even days per domain, depending on the computational resources available. In our method, we save time by only threading against representative templates. Nonetheless, threading is still the major time bottleneck, with a single average-sized (200 residue) domain taking 26 ± 2.5 minutes to thread against the 1,814 templates on one CPU core. To make this more feasible for genome-sized datasets, which typically have thousands or tens of thousands of domains, we have implemented an option for parallel processing of the input sequences. Another possible way to decrease the threading time would be to reduce the number of templates in our library. Preliminary results indicate that, depending on the classifier used, the feature space can be substantially reduced with only a minor impact on classification accuracy. In fact, given our framework, we hypothesize that we can create feature spaces at different scales such that threading can be applied in a hierarchical sequence.

The relationship between the structure of macromolecules to their function is a key annotation principle for computational inference. As the number of solved examples increase, we hypothesize that data-driven feature extraction coupled with machine learning methods as in our method and also in methods like deep learning^29^, will have high utility in extending whole genome/proteome annotations.

## Acknowledgements

This paper was funded in part by DOE CSGF (DE-FG02-97ER25308) to S.A.M. and Health Research Formula Funds from the Pennsylvania Commonwealth to J.K., which disclaims responsibility for any analyses, interpretations or conclusions. We would like to thank Kanishka Rao for contributions to an early version of the feature space.

## Author Contributions

S.A.M. and J.K. conceived the study and wrote the manuscript. S.A.M. implemented the feature space and classifier, applied it to the human proteome, and interpreted results. J.I. contributed to classifier development and validation. All authors read and approved the final manuscript.

## Methods

### Feature extraction and classification

Features were created for each input sequence by constructing empirical coordinate functions comprised of threading scores against a library of 1,814 templates. That is, a given amino-acid sequence was scored against each of these templates by threading to derive 1,814 numerical coordinates. Threading was done using CNFalign_lite from the RaptorX package v.1.62^19^. The templates are the default reference set provided by RaptorX and represent a wide range of different structures with low redundancy, but do not necessarily represent all known folds. We call this feature space the Protein Empirical Structure Space (PESS).

Training sequences were threaded against the templates and the resulting scores were normalized by z-standardization. Test sequences were threaded and normalized using the normalization parameters derived from the training sequences.

We constructed fold predictors over the PESS using both a first Nearest Neighbor (1NN) classifier and Support Vector Machine (SVM) classifier. For the 1NN classifier, pairwise Euclidean distances between each training and testing sequence were calculated, and each test sequence was classified into a fold by finding the closest training neighbor and transferring its fold label to the test sequence. For the support vector machine (SVM) classifier, a linear SVM was trained using the one-vs-all multiclass approach with the C parameter (which controls the penalization of misclassification during training) set to 1/N, where N is the number of positive examples in a given fold.

We also constructed a joint SVM+1NN classifier to assist in identification of fold classes with very small number of training examples. First, a linear SVM was trained as described above to recognize only folds that had at least 20 training examples (“large folds”). The remaining sequences in the training set (“small folds”) were combined into a single class labeled “other”, and this class was not used for classification. A separate 1NN classifier was trained on only the small fold training examples. Classification was then done in two phases: first, all test examples were provided to the SVM, and any test example that received a positive confidence score (based on the signed distance from the hyperplane) was classified into whichever fold gave the highest confidence score; second, the examples that were not classified in the first step were passed to the 1NN model for classification.

All classifiers were implemented in Python using the scikit-learn package^30^.

### Performance assessment

Prediction accuracy was calculated as the fraction of test examples that were classified into the correct fold. Precision (the number of true positives divided by the sum of the true and false positives) and recall (the number of true positive divided by the sum of the true positives and false negatives) were calculated separately for each fold and averaged across the folds. For both precision recall, we excluded cases where the denominator was zero.

### TAXFOLD benchmarks

We obtained three benchmark datasets (EDD, F94, and F195) from the TAXFOLD paper^12^. Each benchmark contains only domain sequences longer than 30 residues with less than 40% pairwise identity, but each contains a different number of folds: EDD contains 3397 sequences in 27 folds, F95 contains 6364 sequences in 95 folds, and F194 contains 8026 sequences in 194 folds. Performance on each dataset was assessed using 10-fold cross validation, with SVM and 1NN classifiers trained and assessed as described above.

### SCOP datasets and final classifier

We downloaded domains from the SCOPe database v2.06 pre-filtered to less than 20% pairwise identity by the Astral database (http://scop.berkeley.edu/astral/ver=2.06), which contained 7,659 domains covering all 1,221 folds in SCOP classes “a” through “g”. We call this dataset “SCOP-20”. We also downloaded the set pre-filtered to 40% identity and removed any domains that were also present in SCOP-20, resulting in 6,322 sequences in 609 folds. We call this dataset “SCOP-40”. We note that almost all SCOP-20 sequences were in SCOP-40 before this filtering, so the final test set has <40% pairwise identity with the training set. We trained a 1NN classifier as described above using the SCOP-20 dataset as training examples and tested the prediction performance using the SCOP-40 set. This classifier was used for all further fold recognition tasks, including the human proteome dataset.

### Human protein analysis

Protein domain sequences for 94 species from the Proteome Folding Project^5^ were downloaded from the Yeast Resource Center public data repository (http://www.yeastrc.org/pdr/pages/download.jsp). To obtain only human sequences, we filtered for protein identifiers marked as “NCBI NR” and had “[Homo sapiens]” in the description. There were a total of 34,330 human domains with length greater than 30 residues, corresponding to 15,619 human proteins.

We classified the domains using the SCOP-20-trained 1NN model with an additional distance threshold to filter out domains that do not belong in any of the represented folds. We determined the threshold nearest-neighbor distance for classification as follows: for each test sequence in SCOP-40, we calculated the nearest neighbor distance before and after removing all SCOP-20 training sequences that belonged to the same fold as the test sequence. We found that a distance threshold of 17.5 provided a good balance between false positives and false negatives (FPR = 9.27%, FNR = 9.49%). After classification with 1NN, only the domains with a nearest-neighbor distance below this threshold we considered confident fold predictions.

Human domain sequences were mapped to PDB entries using a blastp search of PDB requiring that at least 75% of the sequence length had at least 90% identity with a PDB sequence to consider it a match. PDB matches were then mapped to SCOPe classifications using the dir.cla.scope.txt (v.2.06) annotation file downloaded from the SCOPe website.

### RNA-binding proteins

A list of 1,541 known human RBPs was obtained from a recent review^22^. Gene names of the RBPs were matched up to the human protein GIs using the UniProt ID mapping tool, and 1,093 of the RBPs were matched to one or more domains (3,263 domains total). This review also defined a list of 799 Pfam domains with functions related to RNA binding, which we used to filter the 3,263 RBP domains down to those that were most likely to be RNA-binding. Domains were assigned PfamA annotations using hmmscan (http://hmmer.org/). Both a “full-sequence” E ≤ 0.01 and a “best 1” E ≤ 0.1 was required for assignment.

### Novel folds

We extracted all human domains with a nearest neighbor distance ≥30 and performed t-SNE on the PESS projections of these domains using scikit-learn with parameters “perplexity = 10, init = ‘pca’, random_state=123”. Domains were then clustered using DBSCAN from scikit-learn with parameters “eps = 5, min_samples = 5”. Domains and clusters were manually examined for potential boundary prediction errors or previous structural annotations.

### Data and Code Availability

Benchmark datasets, training data, and all human fold and RBP predictions are available at http://kim.bio.upenn.edu/software/pess.shtml. The fold classification source code is freely available at the same website or at https://github.com/sarahmid/PESS.

